# Anatomical mapping and regulation of dopaminergic and octopaminergic neurons during temporal polyethism in the red harvester ant

**DOI:** 10.64898/2026.07.31.742069

**Authors:** Carlos Zavaleta-Zamora, Ingrid Fetter-Pruneda

## Abstract

Many social insects, such as ants, change their behavior stereotypically from nursing to foraging as they age, a concept known as temporal polyethism. Biogenic amines are associated with these specific behaviors, but much about them remains poorly understood in these animals. Many aminergic systems lack anatomical characterization, and little is known about the expression of genes required for the aforementioned behavioral phenotypes. Here, we studied *Pogonomyrmex barbatus* brains and identified the neurons that produce two relevant amines, dopamine and octopamine, by detecting their synthesis enzymes, tyrosine hydroxylase and tyramine beta-hydroxylase, and their transcripts. We also compared the expression of both genes in young nurses and mature foragers by measuring the fluorescence intensity. Dopaminergic and octopaminergic neurons are predominantly located in the protocerebrum and in the subesophageal zone. Neurons that produce dopamine are also present in the optic lobes, whereas octopamine-producing ones show clusters in the antennal lobes. Both genes are downregulated as the organism ages. A reduction in the expression of these genes might be correlated with the mature worker’s increased propensity to perform extranidal tasks, such as foraging.

## Introduction

Division of labor is a hallmark of eusocial insects like ants (Hölldobler and Wilson, 1990; Ward, 2006), with queens being responsible for reproduction, and workers for all other tasks in the colony, such as brood care, nest maintenance, defense, and foraging (Corona et al., 2013; Ingram et al., 2005a; Seid et al., 2005; Seid and Traniello, 2005; Tanaka et al., 2024). Within the worker caste, individuals change their tasks stereotypically as they age, a phenomenon known as temporal polyethism. Young workers predominantly perform intranidal tasks, especially nursing, whereas older workers tend to carry out extranidal duties, including colony defense and foraging (Ingram et al., 2005a; Norman and Hughes, 2016; Seid and Traniello, 2005). This age-related behavioral plasticity provides a powerful framework for linking molecular, neuronal, and behavioral variation within a single genotype.

The molecular and physiological mechanisms underlying division of labor within the worker caste have recently received considerable attention. A variety of molecules have been found to influence particular behaviors in social insects, as in the case of neuropeptides (i.e. inotocin, corazonin) (Fetter-Pruneda et al., 2021; Gospocic et al., 2017) and lipophilic hormones (i.e. juvenile hormone, ecdysteroids) (Norman and Hughes, 2016; Penick et al., 2011; Schulz et al., 2002). Biogenic amines, metabolic derivatives of amino acids, are another relevant group of neuroactive compounds that affect the roles ants take in the colony. These substances strongly influence different processes, including division of labor, queen-like behavior, aggression, and foraging (Friedman et al., 2018; Goolsby et al., 2024; Kamhi and Traniello, 2013; Roeder, 2005; Sasaki et al., 2018; Seid and Traniello, 2005). Additionally, the interaction of two or more of these physiological systems is needed for the occurrence of intricate spatiotemporal phenomena related to complex social behavior, as in the case of juvenile hormone and octopamine (Schulz et al., 2002). Differential gene expression is tightly linked to behavioral heterogeneity seen among castes, and in the task allocation that characterizes worker ants, as in, for example, genes related to vitellogenin (Corona et al., 2013; Tanaka et al., 2024), corazonin (Gospocic et al., 2017), and aminergic synthesis genes (Friedman et al., 2018).

Biogenic amine levels change with age in eusocial insects, and this has been linked with behavioral development (Cuvillier-Hot and Lenoir, 2006; Kamhi and Traniello, 2013; Penick et al., 2011; Seid and Traniello, 2005; Szczuka et al., 2013; Wnuk et al., 2011). In addition, administering drugs that target dopaminergic or octopaminergic systems can alter the likelihood of performing certain tasks or behaviors, such as aggression or foraging (Friedman et al., 2018; Kamhi et al., 2015), mimicking the behavioral changes associated with temporal polyethism.

Despite this extensive functional evidence, the neural substrates through which dopamine and octopamine regulate temporal polyethism remain poorly understood. Precise identification of aminergic neurons is essential for understanding the function of specific neural circuits, and to understand how neuromodulatory systems influence behavioral maturation and task specialization. Moreover, whether age- and task-related behavioral differences are accompanied by changes in the expression of aminergic synthesis genes at the neuronal level is largely unknown. Identifying these neurons and the brain regions they innervate is an essential step toward linking aminergic signaling with the neural circuits underlying age-dependent task specialization.

*P. barbatus* is one of the best studied harvester ant species (Corona et al., 2013; Friedman et al., 2020, 2018; Gordon, 1983; Gordon et al., 2013; Greene and Gordon, 2003; Ingram et al., 2005b; Kamhi et al., 2026), with clear caste differentiation, a monomorphic worker caste, and complex social behaviors such as temporal polyethism. Nevertheless, its aminergic systems have received comparatively less attention at the neuroanatomical level. Transcriptional changes associated with aminergic signaling have been reported, including increased expression of *TBH* in colonies that reduce foraging behavior in response to low environmental humidity as well as changes in foraging behavior in response to dopamine level manipulation (Friedman et al., 2018). Moreover, dopaminergic and octopaminergic systems in ants have not been studied as extensively as in other insects (Evans, 1978; Farooqui, 2012, 2007; Orchard, 1982; Verlinden et al., 2010). Providing foundational neurochemical descriptions is essential for advancing our understanding of ant social behavior and for establishing mechanistic links between environmental cues, aminergic signaling, and colony-level decision-making.

In this study, we map dopaminergic and octopaminergic neurons in the brains of the harvester ant *Pogonomyrmex barbatus* by detecting synthesis enzymes tyrosine hydroxylase (TH) and tyramine beta-hydroxylase (TBH) using immunofluorescence, and their transcripts (*TH, TBH*) using hybridization chain reaction (HCR) RNA fluorescent *in situ* hybridization (RNA-FISH). We further quantify *TH* and *TBH* expression using fluorescence intensity measurements and RT-qPCR in the brains of young nurses and mature foragers. Our results reveal age- and task-associated differences in aminergic gene expression, with a downregulation of both *TH* and *TBH* expression in mature workers. These findings provide a neuroanatomical and molecular framework for understanding how aminergic systems contribute to temporal polyethism in ants.

## Methods

### Ant rearing

Colonies were raised from mated queens captured in 2021 and 2022 in the State of Mexico, Mexico. Two colonies of *Pogonomyrmex barbatus* ants were maintained at a constant temperature of 27 °C (± 1°C), in a 12 h light/12 h dark cycle. The colonies were fed chia seeds, sunflower seeds, canary seed, bee pollen and lyophilized *Tenebrio molitor* larvae every other day.

### Ant selection

The cuticle of adult ants is light-colored at eclosion from the pupa and remains so for about 3 weeks, then darkens as the ants age (Vergara-Martínez et al., 2025). Workers were classified using both age-associated cuticular pigmentation and behavioral context within the colony. Young workers were selected based on their light pigmentation and by their presence inside the nest in close proximity to the brood. Mature workers were selected for their darker pigmentation and for being outside the nest, in the foraging area, near the seed deposits. For the HCR RNA-FISH experiment and brain RNA extraction, we chose eight young and eight mature worker ants from each of two colonies (A and B), for a total of 32 ants. For immunofluorescence, we only chose mature workers.

### TH Immunofluorescence

Ants were anesthetized on ice for 30s, then immersed in ice-cold ethanol (4°C) for 90s to remove cuticular hydrocarbons. Afterward, they were placed in a 1X phosphate-buffered saline (PBS) solution at 4°C under a stereo microscope, and the brains were dissected using ultra-fine forceps. Tracheae and other tissues were discarded, and the brains were immediately immersed in 4% paraformaldehyde (PFA) in cold (4°C) PBS, and fixed for 2 h. All tissues were processed simultaneously. Then, three 5-min washes were performed in PBS at room temperature (25°C). Tissues were then washed and permeabilized in PBS + 0.5% Triton X-100 (PBT) three times for 20 min each, and blocked in PBS + 1% bovine serum albumin (BSA) for 30 min at 25°C. A single wash was performed with 1X PBS + 0.01% Tween 20 (PBTW) for 5 min and incubated in the primary antibody (anti-tyrosine hydroxylase polyclonal, 1:1000, AB152, Sigma-Aldrich, Lot. 3845256, USA) in PBT + 1% BSA overnight at 4°C. Three washes were performed in PBTW for 10 min each at room temperature. The samples were incubated with the secondary antibody (donkey anti-rabbit IgG) labeled with Alexa Fluor 594 (1:500) (Invitrogen, Thermo Fisher Scientific, USA; Ref. A21207), DAPI (4’,6-diamidino-2-phenylindole; Sigma-Aldrich, Israel; Ref. D9542-1MG; Lot. 098M4004V) (1:1000), and phalloidin conjugated to Alexa Fluor 488 (Invitrogen, Thermo Fisher Scientific, USA.; Ref. A12379; Lot. 2486570) (1:400) in PBT for two hours, covered from light. Then, they were washed five times for 10 min each with PBS, and stabilized in the mounting medium VectaShield Plus (Vector Laboratories, USA.; Ref. H-1900; Lot. Z J0401) overnight at 4°C. Finally, they were mounted on slides in the same medium and observed under a microscope.

### TBH Immunofluorescence

TBH antiserum was obtained by rabbit immunization using an ad hoc partial peptide designed from the *Pogonomyrmex barbatus* gene tyramine beta-hydroxylase isoform X2 (XP_011648035.1, LOC105434118, Gene ID: 105434118). The peptide sequence was the following: YDGPCDGADRPEKTQ-amide. Peptide synthesis and antibody production were performed by YenZym Antibodies (California, USA).

Brains were dissected and fixed in glioxal 3% with acetic acid 0.8% in PBS for 1 h, and then processed as above. We incubated the samples with the secondary antibody (donkey anti-rabbit IgG) labeled with Alexa Fluor 488 (1:500) (Invitrogen, Thermo Fisher Scientific, USA; Ref. A21206), DAPI (4’,6-diamidino-2-phenylindole; Sigma-Aldrich, Israel; Ref. D9542-1MG; Lot. 098M4004V) (1:1000), and phalloidin conjugated to Alexa Fluor 647 (Invitrogen, Thermo Fisher Scientific, USA.; Ref. A22287; Lot. 2486570) (1:400) in PBT for two hours, covered from light. After this point the protocol was identical to that for TH Immunofluorescence.

### TH and TBH RNA Fluorescent In Situ Hybridization (FISH)

We used a hybridization chain reaction (HCR, Molecular Instruments) based RNA-FISH assay (Choi et al., 2018; Fetter-Pruneda et al., 2021) to localize the expression of *TH* and *TBH,* genes related to dopamine and octopamine synthetic pathways, respectively. We used RNase AWAY (Invitrogen) and 70% ethanol to clean the tools and workspace to maintain an RNase-free environment and ensure transcript integrity. All reagents used were also RNase-free. Dissections were performed in PBS and immediately transferred to ice-cold PFA (4°C) for at least two hours. The incubation for the following steps (until a different temperature is noted) was done at 4°C. Samples were washed five times for ten minutes each in 0.05% PBT, followed by serial dehydration in ethanol with concentrations of 30%, 50%, 70%, 90%, 99%, 99%, and 99%, with samples left for ten minutes at each step and then left overnight in the last step. The next day, samples were rehydrated in reverse ethanol series (90%, 70%, 50%, 30%) for 10 min at each step, and then incubated in fresh 5% acetic acid (CH_3_COOH) for 5 min. This was followed by five 5-min PBS washes before samples were post-fixed in 2% PFA for 1 h. Afterward, we performed five 10-min washes in 0.05% PBT and then three 10-minute washes in 1% sodium borohydride (NaBH_4_). They were then washed in PBS, five times for 5 min each, and immediately incubated in pre-hybridization buffer (Molecular Instruments, USA; Lot. BPH02923) at 45°C for 30 min. The probe solution was then added, and the samples were incubated overnight at 45°C. This solution contained probes for *P. barbatus TH* (Target: *TH*, Amplifier: B1; Accession number: XM_011633476.1) and *TBH* (Target: *TBH*, Amplifier: B2; Accession number: XM_011649733.2) (Molecular Instruments, USA), at a concentration of 1 pmol (1 μL of 1 μM stock per probe) in 500 μL of HCR probe hybridization buffer (Molecular Instruments, USA). Incubations for the following steps were performed at 45°C. Samples were washed in 100% HCR probe wash buffer (WB) (Molecular Instruments, USA; Lot.BPW03123) for 10 min, then washed serially for 15 min each in 75/25%, 50/50%, and 25/75% WB/5x saline sodium citrate + 0.06% TritonX100 (SSCT), and finally, samples were washed in 100% SSCT for 15 min and then a second time for 30 min. The hairpin solution was prepared (HCR Amplifier B1-h1 + Alexa Fluor 546; Lot. S038723. B1-h2 + Alexa Fluor 546, Lot. S038923; B2-h1 + Alexa Fluor 647, Lot. S039423; B2-h2 + Alexa Fluor 647, Lot. S039723; Molecular Instruments, USA) by the method of snap-cooling: heat to 95°C for 90s and temper for 30 min in a drawer at room temperature. Hairpins were added to the HCR amplification buffer (1:50; Molecular Instruments, USA; Lot. BAM02723), the hairpin solution was added to the tissue, and incubated overnight covered from light at room temperature (25 °C). The next day, five washes of 10 min each were performed in SSCT at room temperature, and the samples were transferred to the mounting medium (SlowFade Diamond, Invitrogen, Thermo Fisher Scientific, USA; Ref. S36963; Lot. 2199092) and incubated overnight at 4°C. The tissues were then mounted on slides in the same mounting medium for observation under the microscope and imaging.

### Confocal Microscopy

Confocal images were acquired with a Nikon A1R+ laser scanning confocal head coupled to an Eclipse Ti-E inverted microscope (Nikon Corporation, Tokyo, Japan) equipped with a motorized stage (TI-S-E, Nikon) and controlled through Nis Elements C v.6.10.02 software located at Unidad de Microscopia IIBO-UNAM, RRID:SCR_022204. Images from the same experiments were taken under identical parameters. To capture the images, we used 20× (Nikon Plan Apo Lambda, NA 0.75), 40× (Nikon Plan Apo Lambda, NA 0.95), or 60× (CFI Plan Apochromat VC 60XC WI, NA 1.20) objectives. Sequential acquisition were done with laser excitation lines of 405, 488, 561, and 640 nm, and light emission detection with standard and GaAsP detectors.

Immunofluorescence images were acquired with a depth of 150 μm, with a step of 3.78 μm for the 20× objective, 0.55 μm for the 40×, and 0.275 μm for the 60×.

For the RNA-FISH experiments, anterior face images were captured at a depth of 150 μm, while those of the posterior face were captured at 100 μm, both with a step size of 1.1 μm, using a 20× objective. Imaging depths were optimized for each orientation based on tissue anatomy and signal quality.

### Image Analysis

Confocal images were analyzed using FIJI software (*Fiji Is Just ImageJ*) (Schindelin et al., 2012). For all images, we used the Despeckle function and 3D Gaussian Blur to generate masks prior to the identification of the bodies of interest. The plug-in “3D Object Counter” was used to detect bodies of interest by thresholding. Finally, integrated density was used to measure fluorescence intensity in the detected objects. Images for figures were modified for clarity using brightness and contrast adjustments, despeckling, background-subtraction filters, and maximum-intensity projections along multiple axes. All images from each experiment were processed with exactly the same adjustment parameters.

### Brain RNA extraction

We used 70% ethanol and RNase AWAY to clean the tools and workspace in order to maintain an RNase-free environment and ensure the integrity of the transcripts. Brain dissections were performed in cold PBS (4°C), and tissue was immediately transferred to 200 μl of cooled (4°C) TRIzol (15596018, Invitrogen). Tissue was quickly homogenized using a pestle, and then, 800 μl TRIzol was added, and mixed by hand. Next, we added 200 μL of chloroform, then mixed the samples vigorously. We centrifuged the tubes for 30 min at 14,000 rpm at 4°C. The aqueous phase was transferred to new tubes, and then we repeated the 200 μl chloroform addition and mixing. Samples were centrifuged again, and we saved the aqueous phase in a new tube with 500 μl cold isopropanol (4°C), which was stored for 2 days at -20°C. Then, the samples were precipitated by centrifuging for 30 min at 14,000 rpm at 4°C. The supernatant was discarded, and the RNA pellet was washed in 75% ethanol. We repeated this centrifugation step twice more. Ethanol was discarded, and the pellets were left to air-dry at room temperature for 3 hours. After that, pellets were resuspended in 20 μl RNase-free ultrapure water.

### Primer design and RT-qPCR

To choose an appropriate reference gene, we tested four housekeeping genes previously reported in another ant species (*Monomorium pharaonis*) *EF1a*, *Act5C*, *TubG2* and *Rp49* (Ding et al., 2022). Primers were designed using Primer-BLAST (Ye et al., 2012), and we subsequently estimated primer quality using the Sequence Manipulation Suite: PCR Primer Stats (Stothard, 2000), except for *Rp49*, which used the same sequence as in Corona et al. (2013). Primers for our genes of interest, *TH* and *TBH*, were designed to match both isoforms reported in NCBI (*TH*: XM_011633476.1, XM_011633477.1; *TBH*: XM_011649732.2, XM_011649733.2) (Table 1).

**Table 1:**
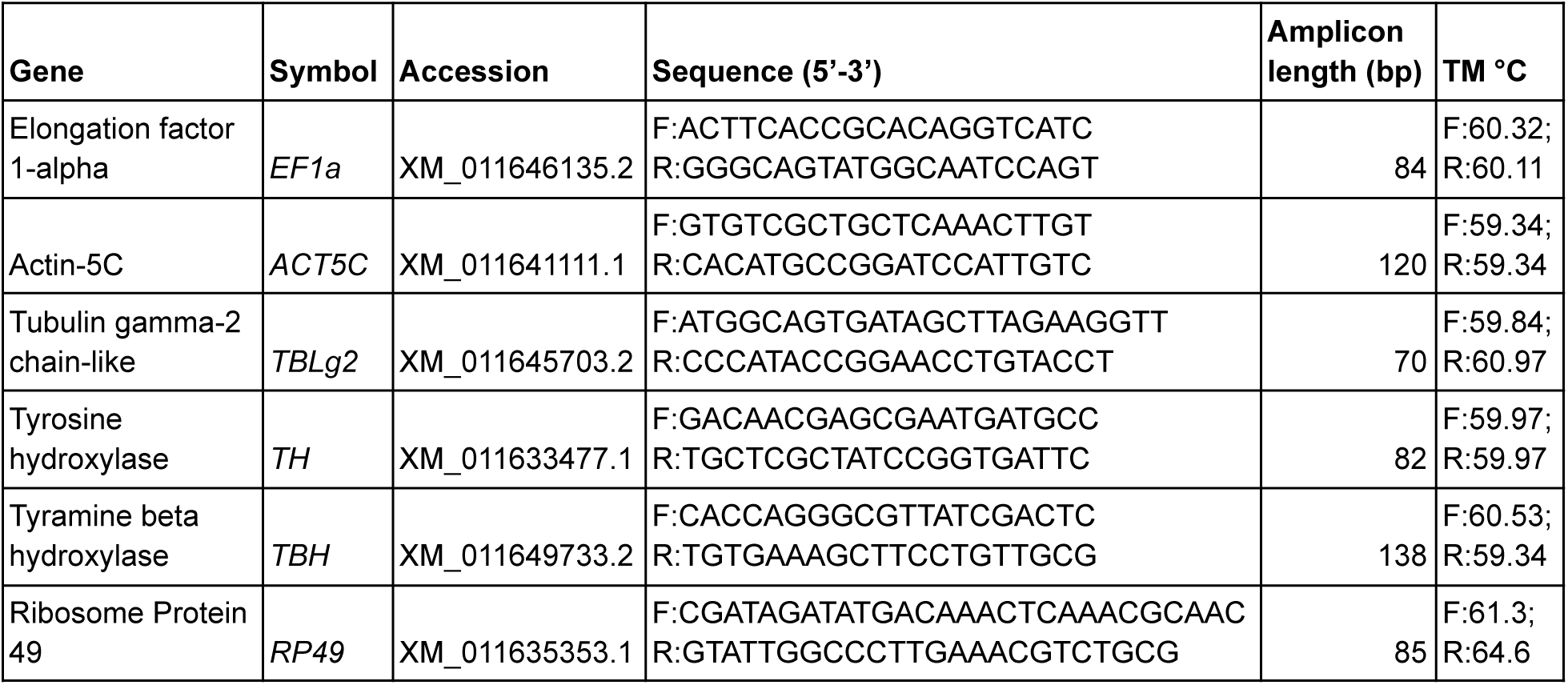
Primer specifications.

Then, we ran an RT-qPCR assay and used four different algorithms to determine the stability in the expression of the candidate reference genes: geNorm, NormFinder, BestKeeper, and dCt (Andersen et al., 2004; Ding et al., 2022; Liu et al., 2022; Pfaffl et al., 2004; Silver et al., 2006; Vandesompele et al., 2002) (Table S1). *Elongation Factor 1 alpha* (*EF1a*) was the best-ranked gene in three of the four algorithms used and was second best in the other and thus was selected for use in further experiments.

We used an RT-qPCR assay to compare *TH* and *TBH* expression between young and mature workers. We chose 16 samples from the RNA extraction based on RNA quality (high concentration and integrity). Three technical replicates were used per sample. For each 96 well plate, we loaded primers of the gene of interest (either *TH* or *TBH*) along with the reference gene *EF1a*.

We used a QuantiNova SYBR Green RT-PCR (Cat. 208154, Lot. 172034682, Qiagen, Germany) kit to perform the experiment. Primer sets were diluted to 10 μM in RNase-free water. RNA samples were diluted 1:10 in 1X QuantiNova Yellow Template Dilution Buffer (Yellow) to get an estimated total of ∼10 ng of RNA per reaction. For each well, we prepared a 10 μL reaction containing 5 μL of 2X QuantiNova SYBR Green RT-PCR Master Mix, 0.1 μL of 100X QuantiNova RT Mix, 1 μL of 10 μM primer solution, 1 μL of RNA template in Yellow, and 2.9 μL of ultrapure water. The exception was the no template control, which did not receive RNA. Then, we ran the experiment in a QIAquant 96 5plex real-time PCR thermocycler (Qiagen, Germany). The program indicated by the kit was used: 10 min at 50 °C, 2 min at 95 °C, and 45 cycles of 5 s at 95 °C and 10 s at 60 °C.

## Results

### Anatomical characterization of TH and TBH positive neurons

To identify dopamine and octopamine-producing neurons, we labeled their synthesis enzymes TH and TBH using immunofluorescence, and their transcripts using HCR RNA-FISH. Both techniques identified most of the same neuronal bodies positive for TH and TBH, respectively (Fig. 1), and anatomical schemes were constructed from these datasets (Fig. 2). The somata of the dopaminergic neurons are distributed in groups in three main neuropils: the protocerebrum, the optic lobes, and the subesophageal zone (SEZ) (Fig. 1a-h). Octopaminergic cell bodies are distributed mainly in the SEZ, the protocerebrum, and the antennal lobes (AL) (Fig. 1i-p). For both aminergic systems (but more evident with TBH), we found generalized staining in the form of punctae in most neuropils, characteristic of neurites.

**Fig. 1.**
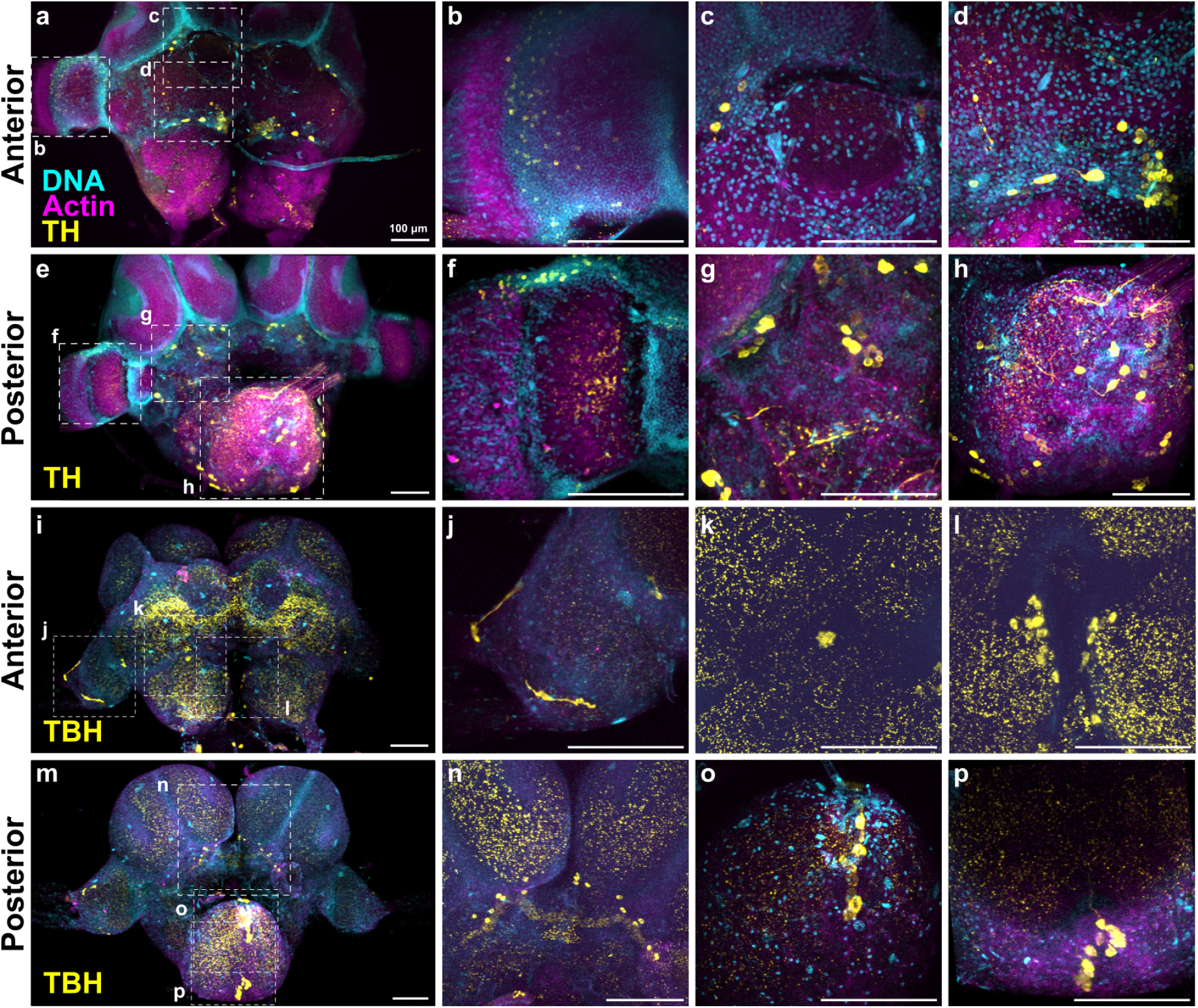
Dopaminergic and octopaminergic cells in the *Pogonomyrmex barbatus* ant brain. Signal shown in yellow indicates TH (a-h) and TBH (i-p) aminergic neurons detected via immunofluorescence. The aminergic somata are predominantly distributed in the protocerebrum (Pr) (c,d,g,n) and subesophageal zone (SEZ) (h,o,p). TH-positive bodies are distributed in the medulla of the optic lobes (b,f). For TBH, somas are found in the antennal lobes (ALs) and Pr border (l). Counterstaining corresponds to DNA (DAPI, cyan) and Actin (phalloidin, magenta). Scale bars correspond to 100 μm.

**Fig. 2.**
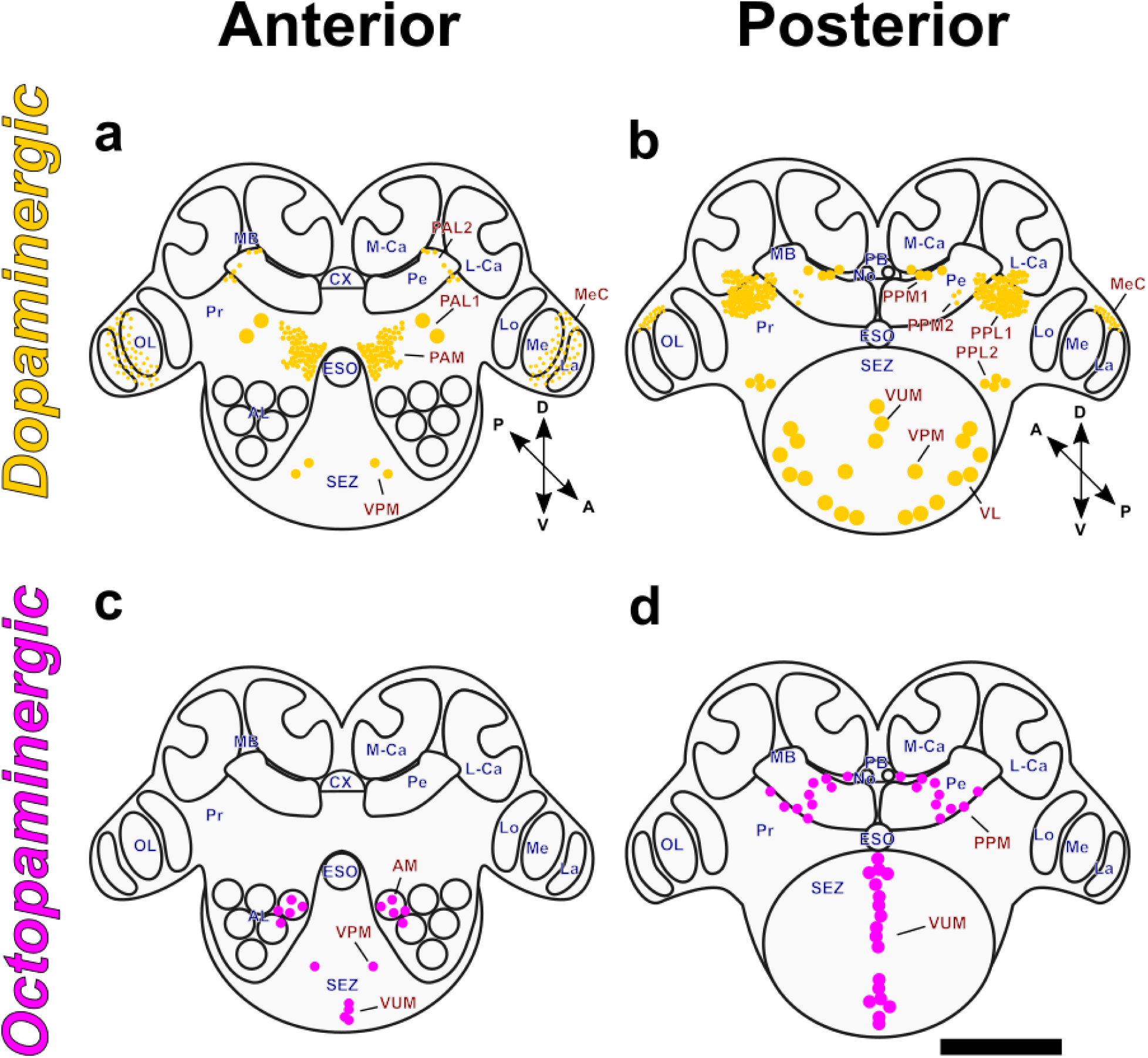
Distribution of dopaminergic and octopaminergic neurons in *P. barbatus* brain, schematic representation. Anterior (a) and posterior (b) views of *TH* and TH distribution in neural tissue. Anterior (c) and posterior (d) views of *TBH* and TBH distribution in neural tissue. Images presented as a stack in the Z axis. Abbreviations: M-Ca, medial calyx of the mushroom bodies; L-Ca, lateral calyx; Pe, peduncle; MB, mushroom bodies; OL, optic lobes; Lo, lobula; medulla; La, lamina; AL, antennal lobes; ESO, esophagus; SEZ, subesophageal zone; AM, antennal medial neurons; MeC, medullar cluster; PAL, protocerebral anterior lateral neurons; PAM, protocerebral anterior medial neurons; VUM, ventral unpaired medial neurons; VPM, ventral paired medial neurons; VL, ventral lateral neurons; PPM, protocerebral posterior medial neurons; PPL, protocerebral posterior lateral neurons. The scale bar corresponds to 200 μm.

The nomenclature used in this work to identify individual neurons or clusters of neurons follows that described by Ito et al. (2014), and other anatomical descriptions in other species, such as *Drosophila melanogaster* (Mao and Davis, 2009; Meissner et al., 2019; Nässel and Elekes, 1992; Sinakevitch and Strausfeld, 2006; White et al., 2010), *Periplaneta americana* (Sinakevitch et al., 2005) and *Apis mellifera* (Sinakevitch et al., 2005; Tedjakumala et al., 2017).

Seven sets of dopaminergic neurons were identified in the protocerebrum, three of which are positioned in the most anterior part of the brain (protocerebral anterior medial [PAM], protocerebral anterior lateral [PAL] 1 and 2) (Fig. 1a) and four in the most posterior (protocerebral posterior medial [PPM] 1 and 2, protocerebral posterior lateral [PPL] 1 and 2) (Fig. 1e). In the optic lobes, we see small positive cells in the medullar region, proximate to lamina (Fig. 1a,b,e,f), that are distributed mainly around the anterior part of the medulla (Fig.1b), but the edges are still perceived in the posterior part (Fig. 1f). In the SEZ (Fig. 1e,h), we observe larger somas than in the protocerebrum, that are distributed in the midline of the SEZ, corresponding to the ventral unpaired medial (VUM) neurons and the ventral paired medial (VPM) neurons. Ventral lateral (VL) neurons are located in the most ventral part of the SEZ and distributed along this neuropil.

In the case of octopaminergic neurons, there are three main regions with cell bodies positive for *TBH* in the anterior view (Fig. 1i). The first one contains the antennal medial (AM) neurons in the peri-esophageal zone of the AL, of which 5-7 cell bodies were positive for *TBH* (Fig. 1l). In the SEZ, we found two VPM and ∼5 VUM *TBH* positive neurons (Fig. 1i). In the immunofluorescence experiment, we found four TBH positive cell bodies that do not match with the signal detected in RNA-FISH, and that also do not share the characteristic shape of other neurons (spheroid with signal absent or dim in the center due to the nucleus). Three are tubular and are located in the OL, two of which are located in the lamina and one near the lobula (Fig. 1i, j,m). The fourth body is located at the edge of the AL (Fig. 1i,k) and apparently lacks a nucleus. In the posterior view, there were 8-10 cell bodies positive for *TBH* on each side of the PPM region (Fig. 1m,n). Near this cluster, we found staining in the central body upper unit and the protocerebral bridge of the central complex (Fig. 1m,n).

In the SEZ, *TBH* positive somata are distributed along the midline, corresponding to the VUM neurons (∼17 cell bodies), divided into superior (Fig. 1m,o) and inferior (Fig. 1 m,p) subgroups.

We found no colocalization of *TH* and *TBH,* suggesting that there are no dual modality dopaminergic-octopaminergic neurons (Fig. S1).

### Age and behavioral caste related changes in *TH* and *TBH* expression

To examine changes in *TH* and *TBH* expression, we compared the fluorescence intensity of young and mature workers detected by HCR RNA-FISH (Fig. 3). We observed an age-related reduction in the number of *TH* and *TBH* transcripts in both anterior and posterior views of the brain (*TH* anterior: Mann-Whitney u = 191.0 p = 0.0011; *TBH* anterior: u = 161.5 p = 0.0416; *TH* posterior: u = 209.0 p = 0.0005; *TBH* posterior: u = 208.0 p = 0.0005) (Fig. 3 e-h), with young workers displaying greater fluorescence intensity than mature workers.

**Fig. 3.**
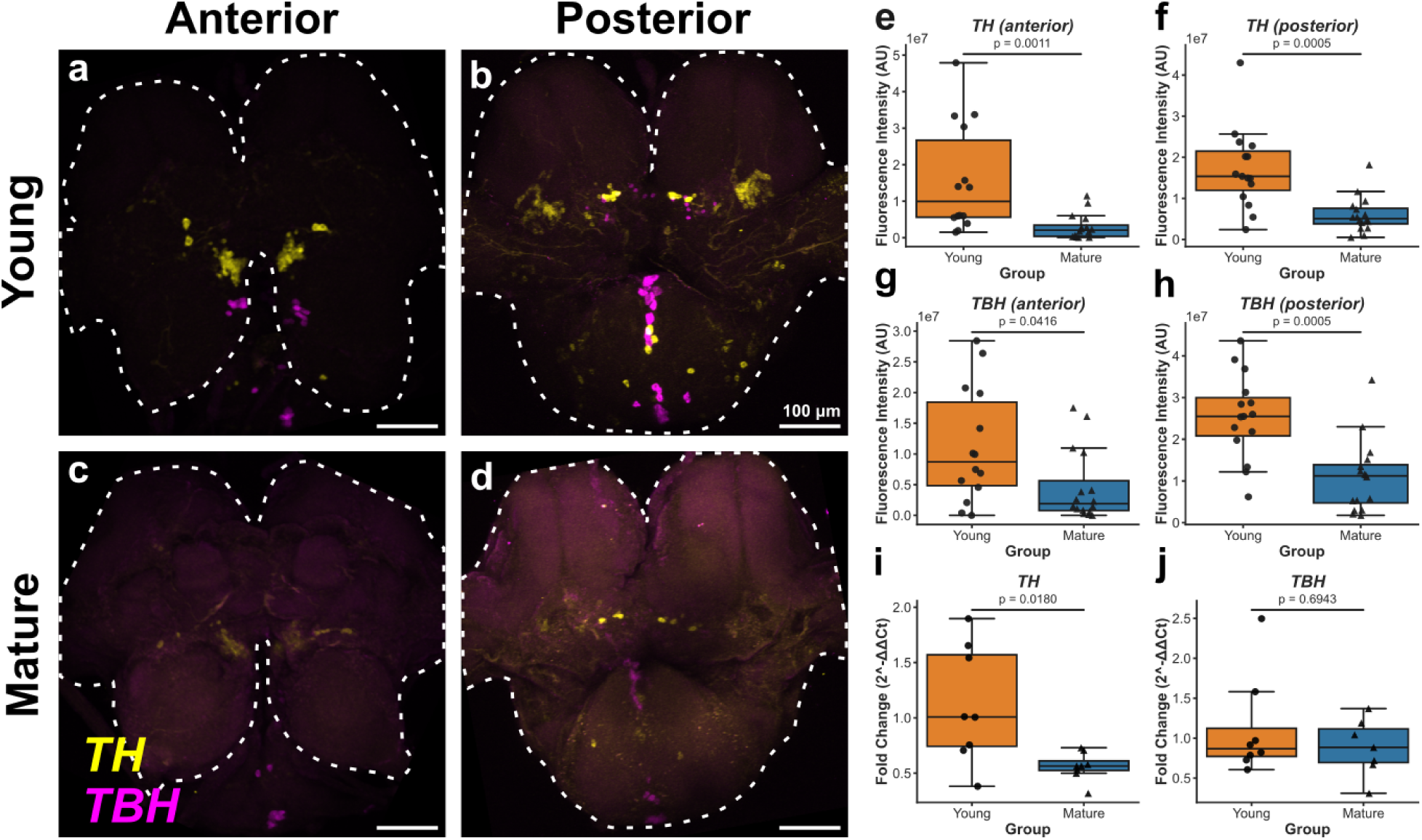
Downregulation of *TH* and *TBH* correlates with age and task. (a-d) Distribution of *TH* (yellow) and *TBH* (magenta) in worker ants’ brains, detected by HCR RNA-FISH in young (a,b) and mature (c,d) individuals. Images correspond to maximum-intensity projections in the Z-axis. Dotted white lines delimit neural tissue. Scale bars correspond to 100 μm. (e-j) *TH* and *TBH* expression quantification using fluorescence intensity analysis (e-h) and RT-qPCR (i,j). Young ants have more *TH* (e,f) and *TBH* (g,h) than mature ants in both the anterior and posterior views. RT-qPCR shows differences only for *TH* (i).

We also performed an RT-qPCR assay and found that young workers showed greater *TH* expression than mature workers (U = 55.0, p = 0.018; Fig. 3 i). We found no significant difference in *TBH* expression between the two groups (U = 32.0, p = 0.694; Fig. 3 j).

## Discussion

In this work, we studied the anatomy of dopaminergic and octopaminergic neurons in the brain of the red harvester ant (*Pogonomyrmex barbatus*) to better understand the effects of aminergic modulation on behavioral shifts intrinsic to temporal polyethism. We also asked how gene expression relates to this behavioral change by studying the expression dynamics of two genes implicated in dopamine and octopamine synthesis, *TH* and *TBH*, respectively, between young nurses and mature foragers.

We found that the distribution of dopaminergic and octopaminergic neurons in ants is similar to that of other insects, present in the protocerebrum, especially near the MBs. Likewise, we see larger dopaminergic and octopaminergic somata in the SEZ, suggesting the anatomical arrangement of these cells could be highly conserved among insects (Fig. 4) (Hoyer et al., 2005; Kamhi et al., 2026; Kreissl et al., 1994; Mao and Davis, 2009; Sinakevitch et al., 2005; Sinakevitch and Strausfeld, 2006; Spivak et al., 2003; Tedjakumala et al., 2017; Zhong et al., 2025).

**Fig. 4.**
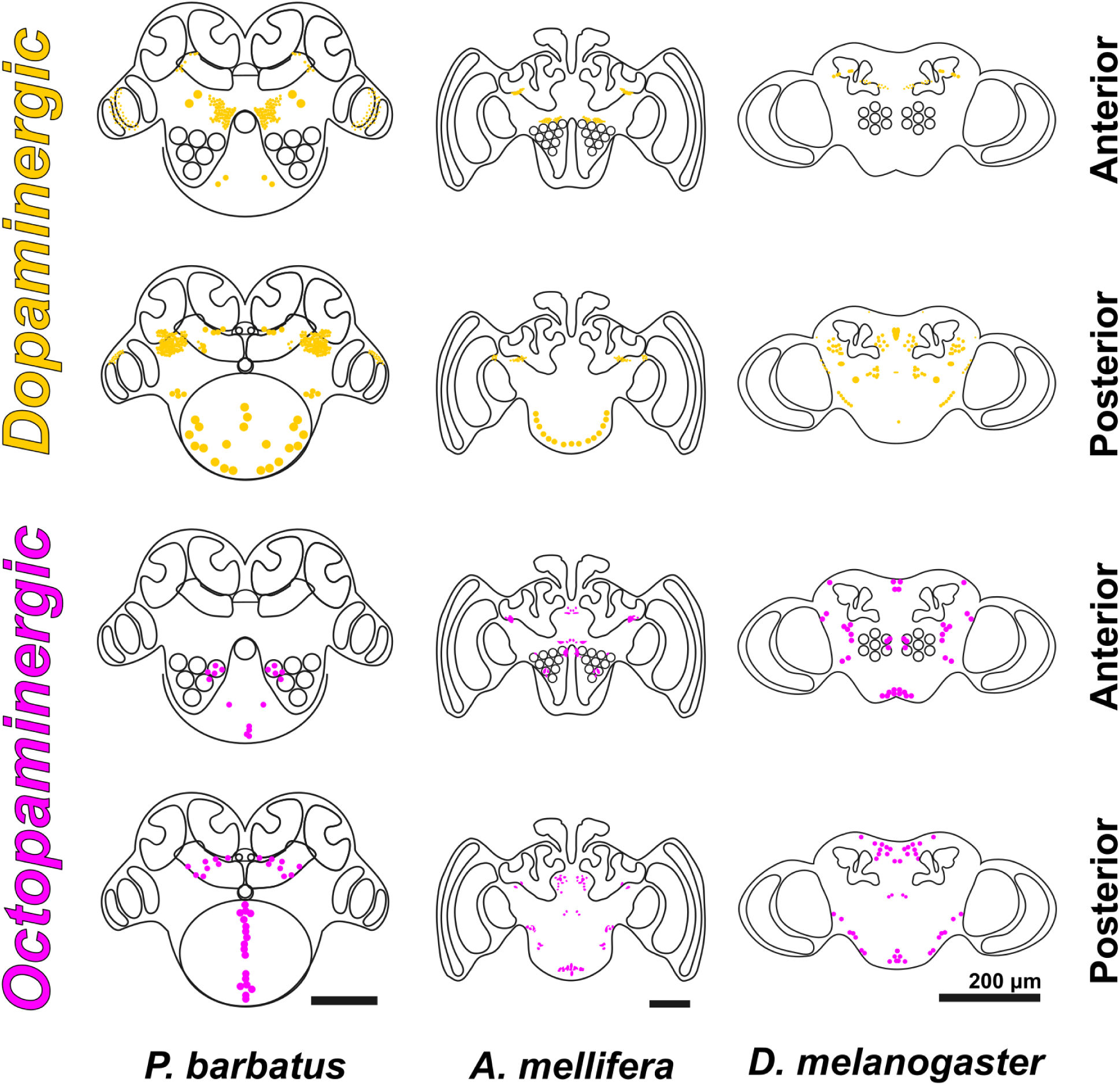
Comparison of the anatomical distribution of aminergic neurons across insect species. Schematic of projections of *P. barbatus, Apis mellifera,* and *Drosophila melanogaster* brains in anterior and posterior view, showing dopaminergic (yellow) and octopaminergic (magenta) cells. Scale bars correspond to 200 μm. The position of the cell bodies in these 2D drawings may not reflect the depth at which they are positioned.

### Characterization of dopaminergic neurons

Dopaminergic cell body distribution is similar to that found in other ants, such as *Monomorium pharaonis* (Zhong et al., 2025). In the anterior view, PAM1 and 2 and PAL TH positive neurons share almost identical distribution and relative size. However, we do not observe a cluster in PAL2 neurons near the OLs in our data. In the posterior view, cluster distribution is almost equivalent between the two species. We lack evidence to further divide the PPM1 neurons into two clusters, as they appear to be only one group. This generates inconsistency in nomenclature, with PPM2 in Zhong et al. (2025) being part of PPM1 in this work and their PPM3 being equivalent to our PPM2, despite the clusters matching in distribution and other characteristics. The same is true for the case of PPL1 neurons, which we cannot split into PPL1 and PPL3 clusters as they have. Further studies are needed to explore the identity of the neurons described above and to determine whether they can be counted as two clusters as in *D. melanogaster* (Mao and Davis, 2009) and pharaoh ants (Zhong et al., 2025).

It has been proposed that PAM and PPL1 neurons are parts of an associative learning circuit, where both are regulated by reward and punishment, allowing flies to display flexible memories and complex learning repertoires (Adel and Griffith, 2021). As in *M. pharaonis* (Zhong et al., 2025), we also found increased extension of these clusters compared to flies, suggesting that the enlargement of these structures could favor complex social behavior and its memory requirements.

Our results show positive dopaminergic structures in the medulla, here named the medullar cluster (MeC). This cluster of neurons coincides with signal in the OL of *P. barbatus* in confocal micrographs recently published by Kamhi et al. (2026), although they are not labeled or discussed in that work. Dopaminergic-related staining in the OLs has been reported before in a variety of insects (Elofsson and Klemm, 1972; Nässel and Elekes, 1992), and somata found in this neuropil correspond to medullar intrinsic neurons (Wendt and Homberg, 1992). The *P. barbatus* MeC identified here might be related to the *D. melanogaster* medullar intrinsic 15 (Mi15) neurons due to similarities in their localization in the OLs and expression of *TH*/*ple*, but they also have some similarity with other medullar neurons such as proximal medullar 3 (Pm3) and Transmedullary 20 (Tm20) neurons (Davis et al., 2020; Meissner et al., 2019). *M. pharaonis* workers lack TH positive cells in the OLs, but staining is noticeable in reproductives, especially in males, which have large eyes and OLs (Zhong et al., 2025). Similarly, no dopaminergic cells were observed in *Ooceraea biroi*, which has significantly reduced OLs (Frank et al., 2025) suggesting that dopaminergic modulation in the MeC may support visual stimulus processing. Dopaminergic neurons were absent in *Harpegnathos saltator*, which is surprising given their large optic lobes and visually oriented behavior. However, the published data excludes parts of the brain where such neurons may be present, leaving their presence or absence unclear (Hoyer et al., 2005). Functional studies will be required to test the role of these cells in visual processing.

We identified a large number of dopaminergic cells in the SEZ, consistent with previous data from *P. barbatus* (Kamhi et al., 2026), and which are also present in other social insects such as *Apis mellifera* and *M. pharaonis* (Tedjakumala et al., 2017; Zhong et al., 2025). As with all other social insects studied, the number of dopaminergic cells identified here surpasses that of flies (Mao and Davis, 2009; Nässel and Elekes, 1992) and locusts (Orchard et al., 1992), suggesting that the precise sensorimotor control needed for complex social behaviors is achieved to some extent through aminergic modulation. Despite their likely relationship to mouthpart sensorimotor processing (Paul and Gronenberg, 2002; Rehder, 1989), and to behavioral specializations, such as nursing and foraging tasks that place very different demands on mouthpart use, the dopaminergic cells of the SEZ have received relatively little attention. Further research is needed to functionally characterize these neurons.

### Characterization of octopaminergic neurons

*P. barbatus* workers exhibit fewer octopaminerigic clusters than other insects, but maintain a similar range of cell counts in clusters that are conserved (10–17 somas in the VUM cluster, 8-10 on each side in the PPM cluster, and ∼10 in the AM cluster) (Busch et al., 2009; Kreissl et al., 1994; Sinakevitch et al., 2005; Sinakevitch and Strausfeld, 2006; Spivak et al., 2003). The neurons in the AM region are consistent with the G3 cluster in *A. mellifera* and the cockroach *Periplaneta americana*, and in the posterior view, we can match PPM neurons to cluster G4, and posterior VUM neurons to cluster 7 (Sinakevitch et al. 2005). The absence of octopaminergic neurons matching the G0 cluster, a protocerebral lateral cluster that supplies the corpora cardiaca through the nervus corporis cardiaci II is notable and raises the question of whether octopamine reaches this neuroendocrine organ by another route in *P. barbatus*. Subsequent experiments will help determine whether the absent clusters were lost in this species’ evolutionary history, are a characteristic specific to the worker caste, or if some other clusters or neurotransmitters have taken over their functions.

Clusters equivalent to the AM cluster in various insects send projections to the OLs (Busch et al., 2009; Sinakevitch et al., 2005; Sinakevitch and Strausfeld, 2006) and in locusts have been related to arousal and attention, specifically with dishabituation of the movement detection system (Busch et al., 2009; Stern, 1999; Stern et al., 1995). The PPM cluster, similar to the G4 cluster, might also innervate parts of the central complex, such as the upper central body and the protocerebral bridge. This suggests that octopaminergic modulation in ants could play an important role in navigation, motor control and visual memory (Homberg et al., 2013). VUM octopaminergic neurons in flies innervate distinct neuropils, such as some regions of the SEZ itself, antennal lobes, protocerebrum, mushroom bodies and central complex (Sinakevitch and Strausfeld, 2006), which might be conserved in *P. barbatus*.

We also observed what appear to be thick TBH positive neuronal fibers in the lamina of the optic lobe, for which we were unable to identify the somas. Characterizing the complete cell morphology and identifying their soma requires additional studies.

### *TH* and *TBH* transcriptional differences between young nurses and mature foragers

Our HCR RNA-FISH experiments show that young intranidal workers expressed higher levels of *TH* and *TBH* transcripts than foragers. This pattern parallels findings in honey bees, where foraging onset is associated with elevated octopamine and serotonin concentrations in the antennal lobes, and where amine-treatment experiments have shown that octopamine specifically, and not serotonin, drives the transition from hive work to foraging (Schulz and Robinson, 2001). Our measurements, however, reflect whole-brain transcript abundance rather than neuropil-specific quantification; we therefore cannot yet say whether the *TH/TBH* decline we observe is distributed evenly across the brain or concentrated in particular regions, such as the antennal lobes. We suggest that both biogenic amines influence the individual probability of performing certain tasks, and that increased expression of both genes promotes a preference for intranidal tasks, including nursing. Whether the age-related decline in expression is a cause or a consequence of task switching, is a distinction that now requires causal manipulation such as pharmacological or RNAi approaches.

This age- and task-based pattern in *P. barbatus* is worth comparing with a recent study in the same species, which provides an anatomical description of the dopaminergic system in *P. barbatus* foragers (Kamhi et al., 2026). Motivated by earlier pharmacological work showing that risk-averse foragers were more sensitive to exogenous dopamine, the authors compared the number of dopaminergic neurons and brain region volume across neuropiles in individuals from colonies with contrasting risk profiles, with respect to water stress. They found no differences in either measurement, and proposed that behavioral differences between colonies instead reflect variation in dopamine synthesis, release, or synaptic distribution rather than in neuron number. Together, these studies suggest that behavioral plasticity in *P. barbatus* is associated primarily with changes in aminergic function rather than gross anatomical remodeling. Whereas Kamhi et al. (2026) found no anatomical differences between colony phenotypes that differ in collective foraging behavior, we detected clear differences in *TH* and *TBH* expression between workers that differ in age and task within a colony. Collectively, these findings point toward regulation of neurotransmitter synthesis and signaling, rather than neuron number, as a major substrate of temporal polyethism. Other studies have described molecular mechanisms associated with behavioral regulation in worker castes, including differential expression of genes such as *forager* (Ingram et al., 2005b), vitellogenins (Corona et al., 2013; Hawkings and Tamborindeguy, 2018; Tanaka et al., 2024), and genes involved in metabolism and signal transduction (Mikheyev and Linksvayer, 2015). This evidence suggests that temporal polyethism is regulated through multiple interacting molecular pathways and highlights the value of integrating complementary approaches to understand its neural basis.

We also quantified *TH* and *TBH* expression by RT-qPCR. While *TH* showed a similar trend with both approaches, *TBH* expression differed between HCR RNA-FISH and RT-qPCR. One possible explanation is that octopaminergic neurons regulate foraging not via transcriptional changes but through differential subcellular trafficking of *TBH* mRNA to neurites, axonal varicosities, or presynaptic boutons, as shown in other transcripts such as *TH* in vertebrates (Gervasi et al., 2016), enabling local synthesis during sustained extranidal activity. If confirmed, this could represent a novel layer of behavioral plasticity in insect aminergic systems. Future studies combining subcellular transcript localization with protein and amine quantification will help clarify how *TBH* expression is regulated during task specialization. Together, our anatomical and transcriptional analyses provide a framework for understanding how aminergic systems contribute to temporal polyethism in ants. Future work integrating neuroanatomy with measurements of neurotransmitter levels, receptor distribution, and functional manipulations will be essential for establishing the causal links between aminergic signaling and task specialization. More broadly, combining anatomical, molecular, and behavioral approaches will be critical for understanding how neural circuits generate complex social behavior.

## Conclusions

This study provides a comprehensive anatomical characterization of the dopaminergic and octopaminergic neuromodulatory systems in the red harvester ant, and provides evidence that changes in *TH* and *TBH* expression are associated with age-related task specialization. Together, these findings establish a neuroanatomical and molecular framework for investigating how aminergic signaling contributes to temporal polyethism and the organization of collective behavior in social insects.

## Acknowledgements

This work was supported by the Global Consortium for Reproductive Longevity and Equality at the Buck Institute for Research on Aging, made possible by the Bia-Echo Foundation (GCRLE-0620 Junior Scholar Award) to I.F.-P. This work was also supported by UNAM-PAPIIT IA206922 & IN210725 funding to I.F-P.

We deeply thank Dr. María Berenice Otero Díaz, and Dr. Jesús Ramírez Santos for their technical assistance. We highly appreciate the technical support provided by Dr. Miguel Tapia Rodríguez from the Unidad de Microscopia IIBO-UNAM. We also thank Dr. Ian A. E. Butler for his helpful comments and edits on the manuscript. Furthermore, we are grateful to all the members of Ant Lab UNAM for their help and comments on the work.

CZZ is a doctoral student of the Programa de Doctorado en Ciencias Bioquímicas at Universidad Nacional Autónoma de México (UNAM) and has received a fellowship (CVU No. 1146622) from Secretaría de Ciencias, Humanidades, Tecnología e Innovación (SECIHTI).

## Notes

### Competing Interest Statement

The authors have declared no competing interest.

