## Supplementary material for "Anatomical mapping and regulation of dopaminergic and octopaminergic neurons during temporal polyethism in the red harvester ant": Figure S1 and Table S1

### Supplementary Information

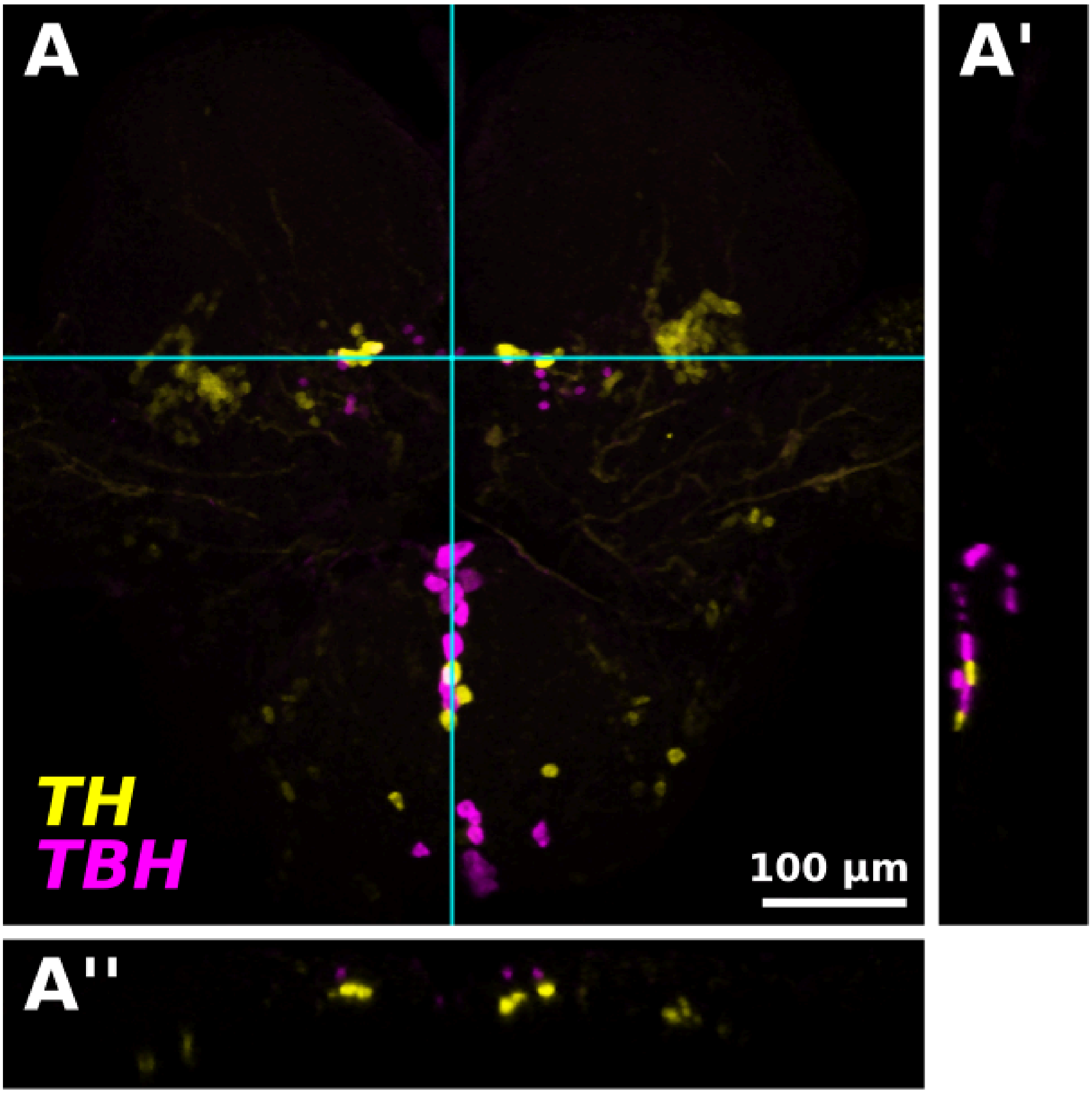


**Fig. S1. Dual-mode dopaminergic-octopaminergic somas are absent in the *P. barbatus* brain.** Distribution of *TH* and *TBH* in worker ants' brains, emphasizing that cell bodies do not coexpress both markers. (A) Projection in the Z axis. (A' and A'') optical slices in the X and Y axes. Cyan lines represent the positions of the optical slices, A' and A'', in A.The scale bar corresponds to 100 µm.

| Gene | geNorm | NormFinder | dCt | BestKeeper (SD) |
|---|---|---|---|---|
| EF1a | 0.461992 | 0.16922 | 0.87882 | 0.388042 |
| RP49 | 0.578081 | 0.58168 | 0.660258 | 0.748396 |
| TUBG2 | 0.542719 | 0.23032 | 1.528675 | 0.429449 |
| ACT5 | 0.530416 | 0.84302 | 1.451735 | 0.843532 |

**Table S1. Expression stability of reference genes.** Expression stability of candidate reference genes using four different algorithms. Lower numbers indicate more stable expression.
